# Non-synaptic alterations in striatal excitability and cholinergic modulation in a SAPAP3 mouse model of compulsive motor behavior

**DOI:** 10.1101/2022.02.07.479446

**Authors:** Jeffrey M. Malgady, Alexander Baez, Kimberly Jimenez, Zachary B. Hobel, Eric M. Prager, Qiangge Zhang, Guoping Feng, Joshua L. Plotkin

**Affiliations:** Department of Neurobiology & Behavior, Stony Brook University Renaissance School of Medicine, Stony Brook, NY 11794; Graduate Program in Neuroscience, College of Arts & Sciences, Stony Brook University, Stony Brook, NY 11794; Medical Scientist Training Program, Renaissance School of Medicine, Stony Brook University, Stony Brook, NY 11794; Center for Nervous System Disorders, Stony Brook University, Stony Brook, NY 11794; McGovern Institute for Brain Research, Department of Brain and Cognitive Sciences, Massachusetts Institute of Technology, Cambridge, MA; Stanley Center for Psychiatric Research, Broad Institute of MIT and Harvard, Cambridge, MA

## Abstract

Deletion of the OCD-associated gene SAP90/PSD-95-associated protein 3 (*Sapap3*), which encodes a postsynaptic anchoring protein at corticostriatal synapses, causes OCD-like motor behaviors in mice. While corticostriatal synaptic dysfunction is central to this phenotype, the striatum efficiently adapts to pathological changes, often in ways that expand upon the original circuit impairment. Here we show that SAPAP3 deletion causes non-synaptic and pathway-specific alterations in dorsolateral striatum circuit function. While somatic excitability was elevated in striatal projection neurons (SPNs), dendritic excitability was exclusively enhanced in direct pathway SPNs. Layered on top of this, cholinergic modulation was altered in opposing ways: striatal cholinergic interneuron density and evoked acetylcholine release were elevated, while basal muscarinic modulation of SPNs was reduced. These data describe how SAPAP3 deletion alters the striatal landscape upon which impaired corticostriatal inputs will act, offering a basis for how pathological synaptic integration and unbalanced striatal output underlying OCD-like behaviors may be shaped.

## Introduction

The compulsive motor symptoms of obsessive-compulsive disorder (OCD) have been attributed to abnormal activity of the cortex-basal ganglia-thalamus-cortex loop (Ahmari and Dougherty, 2015; Burguiere et al., 2015; Graybiel and Rauch, 2000; Pauls et al., 2014; Ting and Feng, 2011). The striatum, the main input nucleus of the basal ganglia, is considered to play an especially central role given that 1) it receives inputs from multiple cortical regions implicated in OCD (Ahmari and Dougherty, 2015), 2) it is overactive in patients (Saxena et al., 1998; Wu et al., 2012), and 3) its normal function involves selecting and setting the urgency of competing motor commands (Gerfen and Surmeier, 2011; Plotkin and Goldberg, 2019; van Maanen et al., 2016), processes that are impaired in OCD. The striatum is composed of two types of striatal projection neurons (SPNs), which provide the structure’s sole output to other nuclei of the basal ganglia: direct pathway SPNs (dSPNs) and indirect pathway SPNs (iSPNs) (Gerfen and Surmeier, 2011; Plotkin and Goldberg, 2019). While action selection involves coordinated activity of both pathways, engagement of dSPNs is associated with action initiation and engagement of iSPNs with action suppression (Cui et al., 2013; Kravitz et al., 2010). Accordingly, circuit models of OCD posit that information flow through the basal ganglia is biased towards the direct pathway (Pittenger et al., 2005; Saxena et al., 1998; Wu et al., 2012).

Mounting evidence suggests that glutamatergic signaling is impaired in OCD (Pittenger, 2015; Pittenger and Bloch, 2014; Rosenberg et al., 2004; Wu et al., 2012). In fact, numerous genes associated with OCD encode proteins that shape postsynaptic glutamate responses, and deletion of these genes in mice induces robust OCD-like motor behaviors such as excessive grooming (Bellini et al., 2018; IOCDF-GC. and OCGAS., 2018; Shmelkov et al., 2010; Welch et al., 2007). Among these OCD-associated genes is SAP90/PSD-95-associated protein 3 (SAPAP3), which encodes a postsynaptic scaffolding protein enriched at corticostriatal synapses (Kindler et al., 2004; Rajendram et al., 2017; Ting and Feng, 2011; Welch et al., 2004; Züchner et al., 2009). SAPAP3 knockout (KO) mice display impaired corticostriatal synaptic function, including reduced AMPA receptor-mediated postsynaptic responses and elevated surface expression of and signaling through metabotropic glutamate receptor 5 (mGluR5) in SPNs (Chen et al., 2011; Wan et al., 2011; Welch et al., 2007). While there is ample evidence for input-specificity of corticostriatal synaptic alterations (Burguiere et al., 2013; Corbit et al., 2019; Hadjas et al., 2020; Wan et al., 2014), evidence for pathway (dSPN vs iSPN) specificity of synaptic impairments is scant (Hadjas et al., 2020). This is especially surprising considering that cortically-driven striatal activity is pathologically biased towards the direct pathway in SAPAP3 KOs (Ade et al., 2016), and pathway-specific optogenetic or chemogenetic manipulation of SPN activity can rescue compulsive grooming behavior in mice lacking either SAPAP3 or its postsynaptic binding partner Shank3 (Ramírez-Armenta et al., 2022; Wang et al., 2017).

Given the extensive overlap of cortical inputs to dSPNs and iSPNs, it is unclear how deletion of SAPAP3 leads to relative changes in striatal output pathway activity. One clue comes from the finding that blocking mGluR5 signaling can “re-balance” the relative cortically-evoked engagement of dSPNs vs iSPNs in SAPAP3 KOs (Ade et al., 2016). Though the mechanism underlying this pathway re-balancing is unclear, given that pathological mGluR5 signaling is observed in both dSPNs and iSPNs (Chen et al., 2011; Wan et al., 2011), it raises the possibility that abnormal glutamatergic signaling induces additional local circuit alterations (Ade et al., 2016). Here we show that constitutive deletion of SAPAP3 leads to intrinsic alterations in SPN excitability and neuromodulation. While the somatic excitability of both dSPNs and iSPNs increased in the dorsolateral striatum of symptomatic mice, dendritic excitability was exclusively enhanced in dSPNs, shifting the normal balance away from the indirect pathway and towards the direct pathway. In addition to changes in intrinsic SPN excitability, we found that SAPAP3 deletion induces a hypercholinergic state in the dorsal striatum, as measured by an increase in striatal cholinergic interneuron (CIN) density and evoked acetylcholine (ACh) release. The elevation in cholinergic tone was paralleled by decreased responsivity of SPNs to basal ACh, in pathway-specific ways. Together, these data suggest that deletion of SAPAP3 induces intrinsic alterations in basal striatal circuitry that will shape how it responds to impaired cortical input.

## Results

### Constitutive deletion of SAPAP3 leads to pathway-specific changes in striatal SPN intrinsic excitability

To determine whether deletion of SAPAP3 leads to non-synaptic pathway-specific alterations in SPNs, we generated a conditional variant of the SAPAP3 knockout mouse (referred to here as SAPAP3 cKI^-/-^) that constitutively lacks SAPAP3 expression and displays robust compulsive grooming behaviors beginning at ~3 months of age (**Figure S1**). We focused our study on striatal output neurons residing in the dorsolateral striatum, the region of the striatum most closely associated with habit learning and habitual behaviors in mice (Balleine and O’Doherty, 2010; O’Hare et al., 2016; Quinn et al., 2013; Yin and Knowlton, 2006; Yin et al., 2004) and known to contain corticostriatal synaptic deficits in SAPAP3 KO mice (Wan et al., 2014; Wan et al., 2011; Welch et al., 2007). To identify direct and indirect pathway SPNs we crossed SAPAP3 cKI^-/-^ mice with either drd1-tdTomato or drd2-eGFP mice (Gong et al., 2003; Shuen et al., 2008) to label dSPNs and iSPNs, respectively. Unlike the central “associative” striatum, where deletion of SAPAP3 has no effect on SPN somatic excitability (Corbit et al., 2019), deletion of SAPAP3 significantly enhanced SPN excitability in the dorsolateral striatum (**Figure 1**). Both dSPNs and iSPNs displayed increased intrinsic somatic excitability, as measured by steeper current-voltage (IV) relationships (**Figure 1A,B**; Mixed-effects analysis; *dSPN* p<0.0001, F(13,344)=13.61, WT n=13, SAPAP3 cKI^-/-^ n=19; *iSPN* p<0.0001, F(13,312)=12.04, WT n=11, SAPAP3 cKI^-/-^ n=20), lower rheobase currents (**Figure 1C**; Mann-Whitney; *dSPN* p=0.0014, WT n=15, SAPAP3 cKI^-/-^ n=21; *iSPN* p=0.0024, WT n=13, SAPAP3 cKI^-/-^ n=24) and higher input resistances (**Figure 1D**; Mann-Whitney; *dSPN* p<0.0001, WT n=15, SAPAP3 cKI^-/-^ n=21; *iSPN* p=0.0007, WT n=13, SAPAP3 cKI^-/-^ n=21). In addition, dSPNs also exhibited a more depolarized resting membrane potential, adding a layer of pathway specificity to the observed changes in somatic excitability (**Figure 1E**; Mann-Whitney; *dSPN* p=0.0001, WT n=16, SAPAP3 cKI^-/-^ n=21; *iSPN* p=0.2665, WT n=21, SAPAP3 cKI^-/-^ n=27). Such changes are consistent with increased firing rates observed in SAPAP3 KOs *in vivo* (Burguiere et al., 2013).

**Figure 1:**
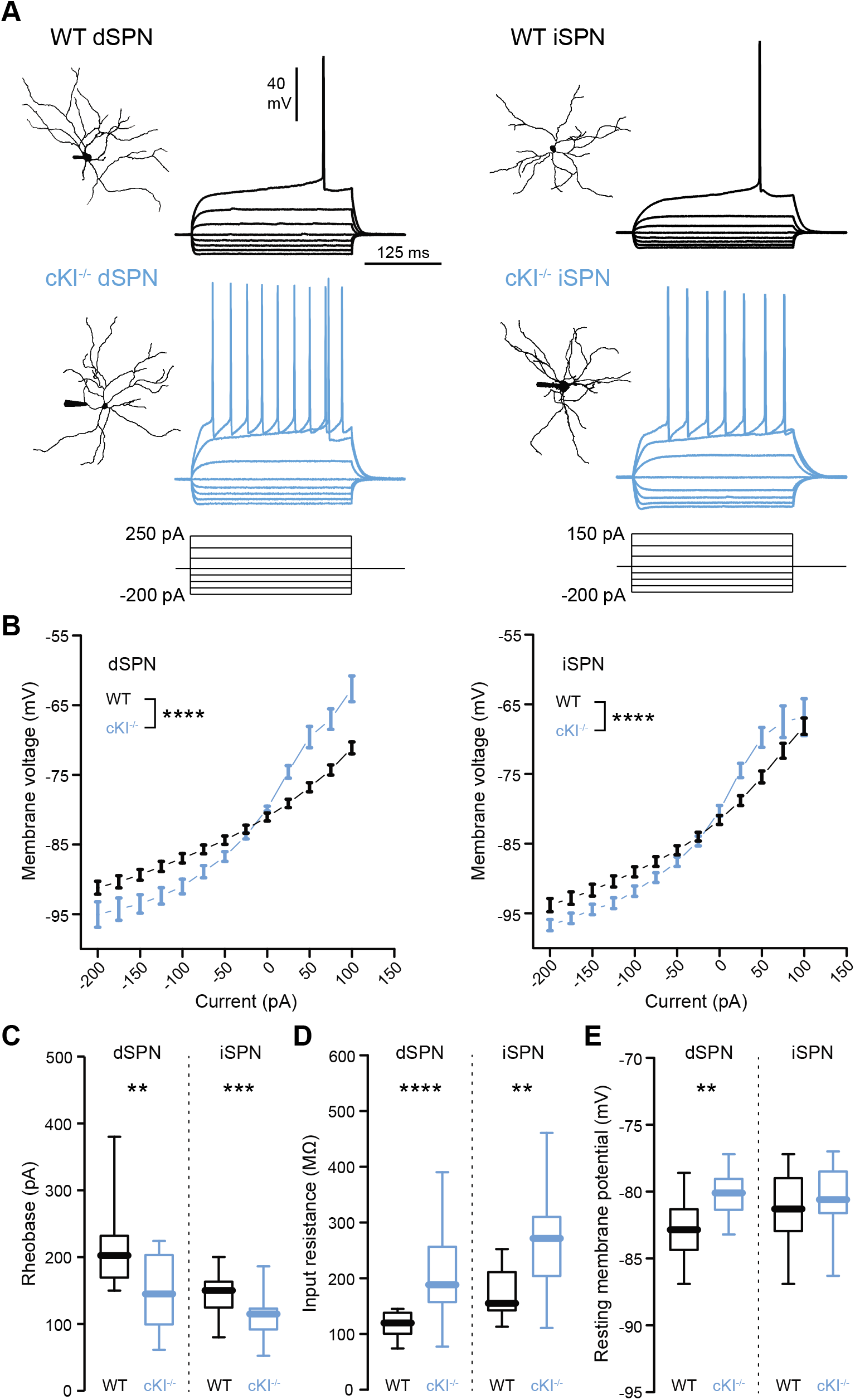
Increased somatic excitability of dSPNs and iSPNs in the dorsolateral striatum of SAPAP3 cKI^-/-^ mice. (A) Example reconstructions & current-voltage (IV) traces of WT (*top*; black) and cKI^-/-^ (*bottom;* blue) dSPNs (*left*) and iSPNs (*right*). (B) Quantification of steady state voltage in response to increasing current injections in WT and cKI^-/-^ dSPNs (*left*) and iSPNs (*right*). (C) Box plots showing rheobase currents in WT and cKI^-/-^ SPNs. *p<0.05, **p<0.01, ***p<0.001, ****p<0.0001. (D) Box plots showing input resistance in WT and cKI^-/-^ SPNs. *p<0.05, **p<0.01, ***p<0.001, ****p<0.0001. (E) Box plots showing resting membrane potential in WT and cKI^-/-^ SPNs. *p<0.05, **p<0.01, ***p<0.001, ****p<0.0001.

While deletion of SAPAP3 increased the excitability of SPN soma, it is their dendrites that contain the SAPAP3-enriched postsynaptic densities that are targeted by corticostriatal synapses (Wan et al., 2014; Welch et al., 2007) and directly shape synaptic integration (Carter and Sabatini, 2004; Plotkin et al., 2011; Plotkin et al., 2013). We measured the degradation of backpropagating action potential (bAP)-evoked local Ca^2+^ transients, from proximal to distal dendrites, as a surrogate marker of dendritic excitability in SAPAP3 cKI^-/-^ mice (Carrillo-Reid et al., 2019; Day et al., 2008) (**Figure 2A**). bAP-evoked Ca^2+^ transients were significantly larger in the distal dendrites of dSPNs in SAPAP3 cKI^-/-^ mice, suggesting enhanced dendritic excitability (**Figure 2B,C;** Mann-Whitney; p=0.021, WT n=31, SAPAP3 cKI^-/-^ n=31). The dendritic excitability of iSPNs, however, was unaltered (**Figure 2B,C**; Mann-Whitney; p=0.7424, WT n=30, SAPAP3 cKI^-/-^ n=32). This pathological enhancement of dSPN dendritic excitability (and lack of change in iSPNs) was also observed in response to sustained bursts of bAPs mimicking ongoing activity (**Figure 2D,E**; Mann-Whitney; *dSPN* p=0.0036, WT n=34, SAPAP3 cKI^-/-^ n=42; *iSPN* p=0.7488, WT n=29, SAPAP3 cKI^-/-^ n=39). Thus, constitutive deletion of SAPAP3 triggers pathway-specific adaptations in the intrinsic excitability of SPNs that are consistent with biased striatal output towards the direct pathway.

**Figure 2:**
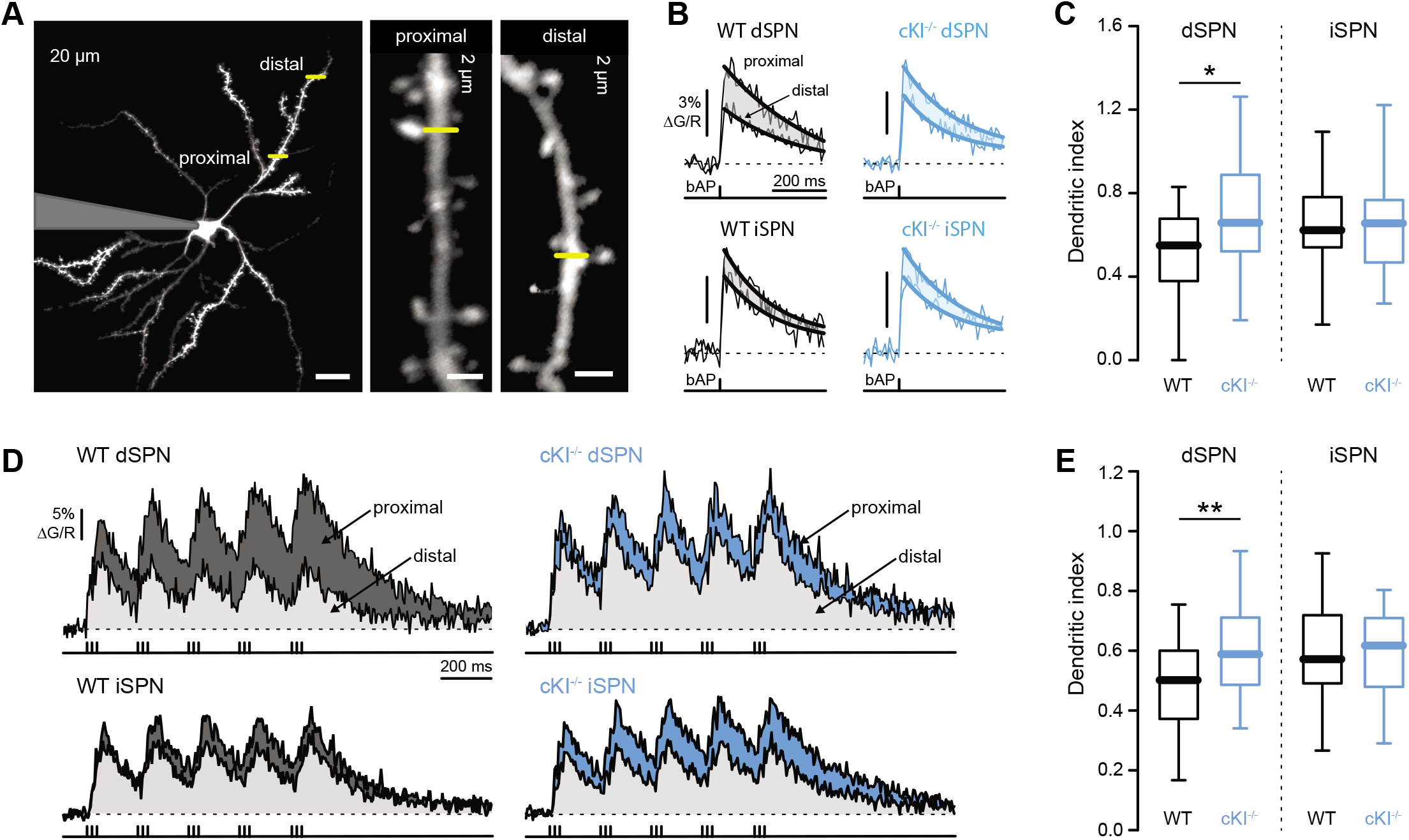
Dendritic excitability of dSPNs is increased in the dorsolateral striatum of SAPAP3 cKI^-/-^ mice. (A) 2-photon maximum intensity projection of a dSPN from a SAPAP3 cKI^-/-^ x drd1-tdTomato mouse (left); high mag images of the proximal and distal dendritic regions (right). Yellow bars indicate sites of Ca^2+^ line scans. Scale bars = 15 μm (left) and 2 μm (right). (B) Averaged proximal & distal bAP-evoke Ca^2+^ transients with overlaid exponential fits (difference shaded) from dendritic shafts of WT (*left;* black) and SAPAP3 cKI^-/-^ (*right;* blue) dSPNs (*top*) and iSPNs (*bottom*). Stimulus protocol eliciting bAP shown time-locked below traces. (C) Box plots of SPN dendritic indexes (ratio of distal:proximal bAP-evoked Ca^2+^ transients) evoked by single bAPs in WT and SAPAP3 cKI^-/-^ mice. *p<0.05, **p<0.01. (D) Averaged proximal (dark gray) & distal (light gray) dendritic shaft Ca^2+^ transients recorded from WT (*left;* black) and SAPAP3 cKI^-/-^ (*right*; blue) dSPNs (*top*) and iSPNs (*bottom*) in response to bursts of bAPs. Stimulus protocol eliciting bAPs shown time-locked below traces. (E) Box plots of SPN dendritic indexes in response to bursts of bAPs in WT and SAPAP3 cKI^-/-^ mice. **p<0.01.

### Constitutive deletion of SAPAP3 generates a hypercholinergic state in the dorsal striatum

While striatal output is driven by synaptic inputs to SPNs, it is profoundly shaped by non-synaptic processes such as the intrinsic excitabilities of SPNs (above) and intrastriatal release of neuromodulators (Gerfen and Surmeier, 2011; Plotkin and Goldberg, 2019). Among the most influential of these neuromodulators are monoamines (such as dopamine and serotonin) and ACh. There is evidence for abnormal monoaminergic signaling in both OCD patients and animal models of OCD (Bellini et al., 2018; Fineberg et al., 2011; Pauls et al., 2014; Wood et al., 2018), and experimental manipulation of striatal ACh can induce tic- and OCD-like behavioral stereotypies in rodents (Aliane et al., 2011; Crittenden et al., 2014; Xu et al., 2015). While the sources of monoamines are extrastriatal, the predominant source of striatal ACh is from cholinergic interneurons (CINs) residing within the striatum itself (Gerfen and Surmeier, 2011; Plotkin and Goldberg, 2019; Zhou et al., 2002). We therefore investigated the impact of constitutive SAPAP3 deletion on striatal cholinergic signaling. Choline Acetyltransferase (ChAT) immunohistochemistry was performed to visualize and quantify striatal CIN density in the precommissural striatum of WT and symptomatic SAPAP3 cKI^-/-^ mice (**Figure 3**). Strikingly, we found that striatal CIN density was increased in SAPAP3 cKI^-/-^ mice (**Figure 3A,B**; Mann-Whitney; p=0.0048; WT n=10, SAPAP3 cKI^-/-^ n=11). The increase in CIN density was restricted to the dorsal striatum (**Figure 3B, Figure S2A**; Mann-Whitney; *dorsal* p=0.0021; *ventral* p=0.2512; WT n=10, SAPAP3 cKI^-/-^ n=11), and displayed a slight caudal to rostral gradient (**Figure 3C**; Mixed effects analysis; p=0.0033, F(1,19)=11.27; WT n=10, SAPAP3 cKI^-/-^ n=11). No difference was observed in striatal volume (data not shown).

**Figure 3:**
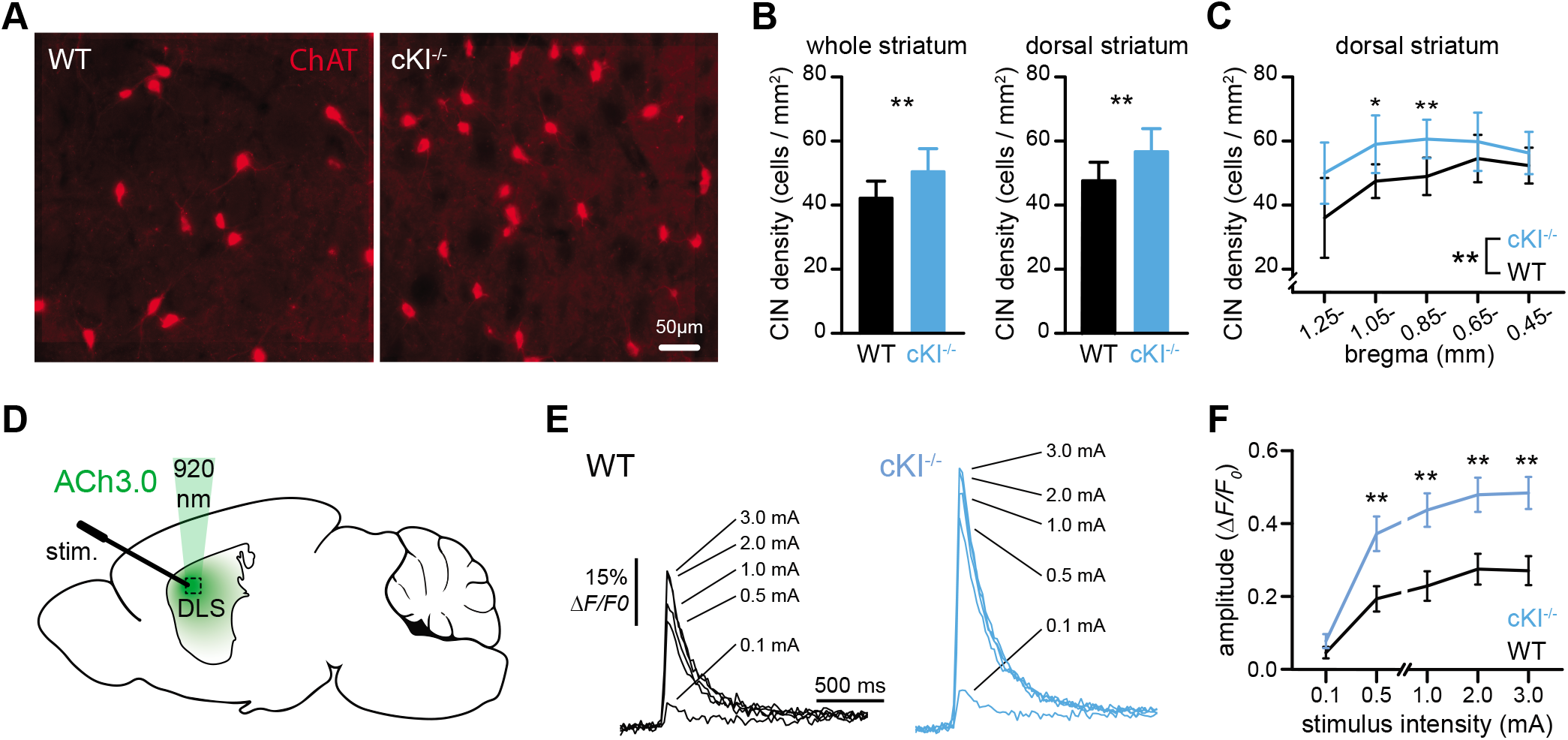
CIN density and evoked ACh release are increased in the dorsal striatum of SAPAP3 cKI^-/-^ mice. (A) Representative images showing choline acetyltransferase (ChAT) staining in the dorsolateral striatum of WT (*left*) and SAPAP3 cKI^-/-^ (*right*) mice. (B) Bar graphs showing CIN densities in the entire striatum (*left*) and dorsal striatum (*right*) of WT (black) and SAPAP3 cKI^-/-^ (blue) mice. (C) Line graphs showing CIN densities in the dorsal striatum of WT (black) and SAPAP3 cKI^-/-^ (blue) mice binned rostro-caudally from bregma 1.25-0.25mm. (D) Schematic of experimental design used in (E and F). (E) Averaged line scans measuring changes in ACh3.0 fluorescence in dorsal striatum neuropil of WT (black) and SAPAP3 cKI^-/-^ (blue) mice in response to increasing electrical stimuli. (F) Line graphs showing evoked ACh3.0 fluorescence amplitudes (calculated from exponential fits) at increasing stimulus intensities in WT (black) and SAPAP3 cKI^-/-^ (blue) mice. **p<0.01

Does the increase in CIN density in SAPAP3 cKI^-/-^ mice translate to an increase in ACh release? To test this, we infected the dorsal striatum with the genetically encoded ACh sensor GRAB_ACh3.0_ (ACh3.0) (Jing et al., 2020) and used 2-photon laser scanning microscopy (2PLSM) to measure electrically evoked ACh release in acute brain slices (**Figure 3D**). Electrically-evoked ACh3.0 fluorescence signals were significantly larger in SAPAP3 cKI^-/-^ mice across a large range of stimulus intensities (**Figure 3E,F**; Mann-Whitney; 0.1mA p=0.1716, 0.5mA p=0.0067, 1.0mA p=0.0019, 2.0mA p=0.0048, 3.0mA p=0.0017; WT n=17, 2 mice, SAPAP3 cKI^-/-^ n=19, 3 mice). Specificity of the fluorescence signal as an indicator of sensor-ACh binding was confirmed using the muscarinic ACh receptor antagonist scopolamine hydrobromide (10 μM) (**Figure S2B,C**).

### Constitutive deletion of SAPAP3 attenuates cholinergic signaling in striatal SPNs

Although striatal levels of ACh may be higher in SAPAP3 cKI^-/-^ mice, it is not clear how SPNs perceive such an increase. SPNs express types 1 and 4 muscarinic receptors (M1 and M4, respectively), but not nicotinic receptors (Assous, 2021; Goldberg et al., 2012; Shen et al., 2015). Muscarinic receptors modulate a constellation of conductances that ultimately regulate SPN somatic and dendritic excitability (Akins et al., 1990; Day et al., 2008; Goldberg et al., 2012; Shen et al., 2007). While a global increase in muscarinic signaling itself would not readily explain the dSPN-specific *enhancement* in dendritic excitability described above, alterations in the responses to ACh via muscarinic receptors may provide further insight. We therefore measured SPN dendritic excitability in *ex vivo* slices from WT and SAPAP3 cKI^-/-^ mice before and after bath application of the muscarinic antagonist atropine (10 μM); CINs were spontaneously active in slices from both genotypes under these conditions (data not shown). Atropine reduced dendritic excitability in both dSPNs and iSPNs from WT mice, but not SAPAP3 cKI^-/-^ mice (**Figure 4A,B**; B: pre vs post atropine: Wilcoxon matched pairs; WT dSPN p=0.0305, n=17, SAPAP3 cKI^-/-^ dSPN p=0.2734, n=13, WT iSPN p=0.0105, n=13, SAPAP3 cKI^-/-^ iSPN p=0.2069, n=17; WT vs SAPAP3 cKI^-/-^: Mann-Whitney; dSPN p=0.016, iSPN p=0.0512).

**Figure 4:**
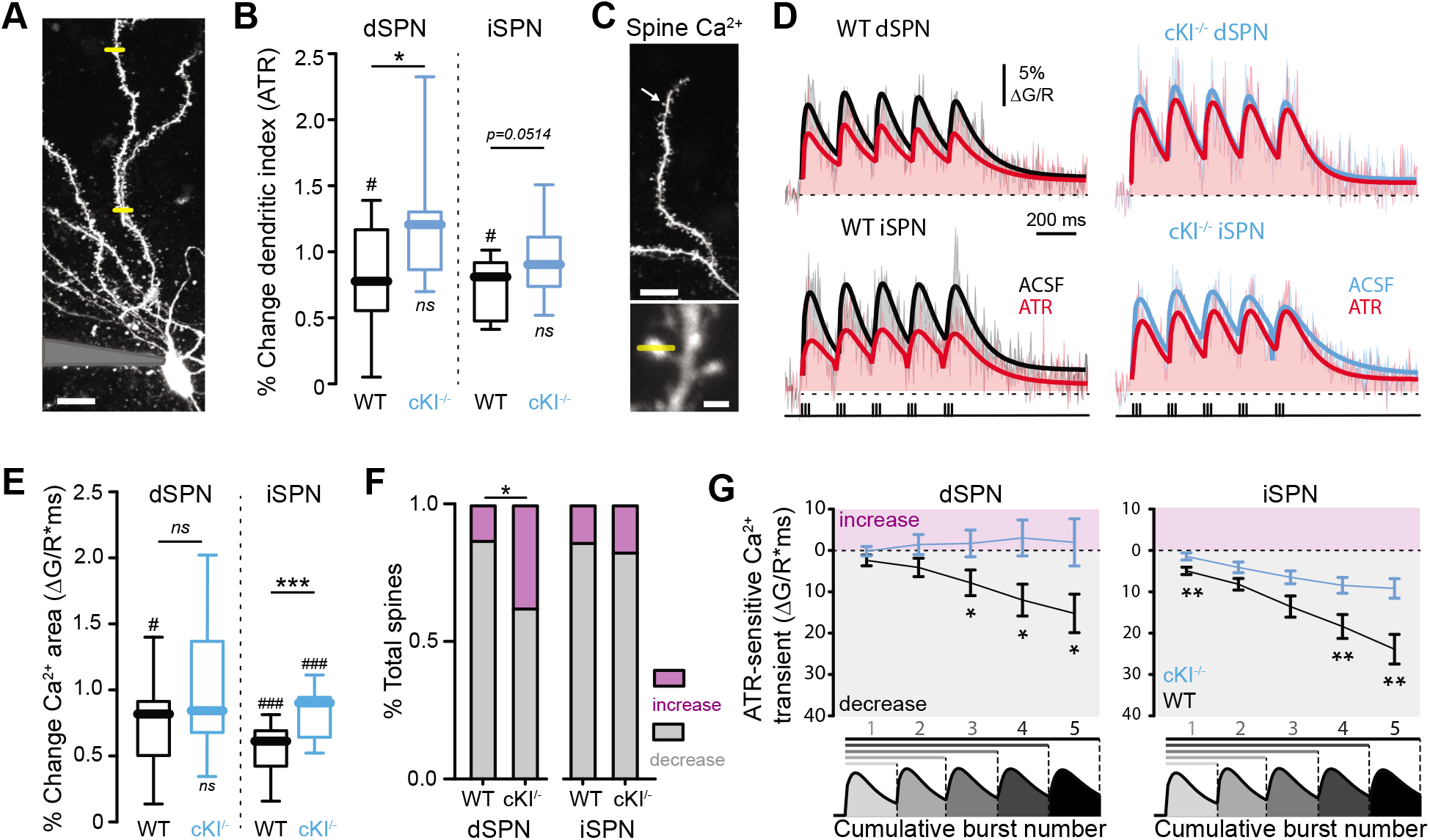
Muscarinic signaling is impaired in dSPNs and iSPNs of SAPAP3 cKI^-/-^ mice. (A) 2-photon maximum intensity projection of a dSPN; sites of proximal and distal Ca^2+^ line scans are shown (yellow bars). Scale bar = 15 μm. (B) Box plots to the right show changes in SPN dendritic indexes (measured in response to bursts of bAPs) induced by atropine (10 μM) in WT and SAPAP3 cKI^-/-^ mice. # indicates a statistically significant difference between pre-post atropine application; * indicates a statistically significant difference between genotypes. #p<0.05, *p<0.05, ns-not significant. (C) 2-photon maximum intensity projection of a dSPN; arrow indicates location of high mag image shown below (top). High mag image of a dendritic spine and corresponding Ca^2+^ line scan (bottom). Scale bars = 15 μm (top) and 1 μm (bottom). (D) Averaged Ca^2+^ transients (evoked by bursts of bAPs) recorded from distal dendritic spines of WT (*top*; black) and SAPAP3 cKI^-/-^ (*bottom*; blue) dSPNs (*left*) and iSPNs (*right*) before (ACSF, black) and after bath application of 10μM atropine (ATR; red). Stimulus protocol eliciting bAPs shown time-locked below traces. (E) Box plots showing atropine-induced changes in total Ca^2+^ transients (evoked by bursts of bAPs) in distal dendritic spines of WT and SAPAP3 cKI^-/-^ SPNs. #p<0.05, *p<0.05, ##p<0.01, **p<0.01, ###p<0.001, ***p<0.001, ns-not significant. (F) Bar graphs showing fractions of total imaged spines whose bAP burst-evoked Ca^2+^ transients were decreased (gray) or increased (magenta) by atropine. *p<0.05 (G) Line graphs showing the cumulative atropine-sensitive components of dendritic spine Ca^2+^ transients evoked by 5 successive bursts of bAPs; Ca^2+^ transients were divided into segments corresponding to each of the 5 successive bursts (diagrammatically shown in register). Y-axis is reported as an absolute value of positive areas (indicating a net atropine-induced decrease; gray) and negative areas (indicating a net atropine-induced increase; magenta). *p<0.05, **p<0.01.

In the above experiments, dendritic excitability was estimated from the relative decrement of bAP-evoked Ca^2+^ transients measured along SPN dendritic *shafts*. In addition to somatic excitability and bAP propagation, muscarinic receptors modulate local Ca^2+^ influx and mobilization from intracellular stores (Howe and Surmeier, 1995; Oldenburg and Ding, 2011; Olson et al., 2005; Pisani et al., 2007). We therefore measured the modulatory effect of atropine on evoked Ca^2+^ transients within SPN distal dendritic spines, as an additional readout of muscarinic signaling (**Figure 4C,D**). Atropine significantly reduced spinous Ca^2+^ transients evoked by bursts of bAPs in both dSPNs and iSPNs from WT mice. This modulation, however, was significantly attenuated in SPNs from SAPAP3 cKI^-/-^ mice (**Figure 4E;** pre vs post atropine: Wilcoxon matched pairs; WT dSPN p=0.0208, n=18, SAPAP3 cKI^-/-^ dSPN p>0.9999, n=15, WT iSPN p=0.0002, n=14, SAPAP3 cKI^-/-^ iSPN p=0.0001, n=20; WT vs SAPAP3 cKI^-/-^: Mann-Whitney; dSPN p=0.3376, iSPN p=0.0009). While muscarinic signaling was reduced in both SPN types in SAPAP3 cKI^-/-^ mice, this occurred in different ways. In iSPNs the overall percent change in Ca^2+^ transients induced by atropine was reduced across the population of measured spines in the mutants. In dSPNs, however, the proportion of spines in which atropine increased vs decreased Ca^2+^ transients was enlarged (**Figure 4F**; Fisher’s exact; dSPN p=0.0332, WT n=31 spines, SAPAP3 cKI^-/-^ n=24 spines; iSPN p=0.7238, WT n=22 spines, SAPAP3 cKI^-/-^ n=35). Further analysis of the atropine-sensitive component of spine Ca^2+^ transients revealed an activity-dependent aspect of muscarinic modulation: both normal muscarinic signaling and impairment of muscarinic signaling in SAPAP3 cKI^-/-^ mice become more prominent as spiking activity continues (**Figure 4G**; Mann-Whitney; dSPN WT n=18, SAPAP3 cKI^-/-^ n=16; iSPN WT n=14, SAPAP3 cKI^-/-^ n=20), suggesting that deletion of SAPAP3 may impair the striatum’s ability to effectively modulate extended bouts of striatal activity.

## Discussion

This study reveals that deletion of the OCD-associated synaptic gene SAPAP3, which causes corticostriatal synaptic dysfunctions and compulsive overgrooming, results in broad non-synaptic alterations in striatal neuronal excitability and neuromodulation. This includes a general increase in SPN somatic excitability, pathway-specific augmentation of dSPN dendritic excitability, hypercholinergic tone and impaired muscarinic signaling. Together, these data describe how manipulation of a single postsynaptic risk gene can lead to widespread alterations in intrinsic striatal function that will, ultimately, change the landscape upon which impaired corticostriatal synapses act. It also highlights the role of the cholinergic system as a potential target for therapeutic intervention in OCD.

We restricted our study to the dorsolateral striatum for several reasons: 1) it has been proposed that compulsions may represent exaggerated habits (Burguiere et al., 2015; Graybiel and Rauch, 2000), 2) habit learning is dysregulated in SAPAP3 KO mice (Ehmer et al., 2020; Hadjas et al., 2019), 3) habit learning has been linked to activity in the dorsolateral striatum of mice (Balleine and O’Doherty, 2010; O’Hare et al., 2016; Quinn et al., 2013; Vicente et al., 2016; Yin and Knowlton, 2006; Yin et al., 2004) and 4) corticostriatal synapses within the dorsolateral striatum are impaired in SAPAP3 KO mice (Wan et al., 2014; Wan et al., 2011; Welch et al., 2007). Unlike the central “associative” striatum of SAPAP3 KO mice (which also displays corticostriatal synaptic impairments) (Corbit et al., 2019), we found that somatic excitability is elevated in both direct and indirect pathway SPNs in the dorsolateral striatum. This is consistent with observations that baseline firing rates and activity are increased in the striatum of SAPAP3 KOs (Burguiere et al., 2013; Mintzopoulos et al., 2016) and striatal activity is elevated in OCD patients (Saxena et al., 1998). In contrast to somatic excitability, dendritic excitability was exclusively enhanced in dSPNs. As all cortical inputs to SPNs target dendrites (Gerfen and Surmeier, 2011), and dendritic excitability profoundly shapes how SPNs respond to convergent corticostriatal inputs (Plotkin et al., 2011; Prager et al., 2020), this pathway-specific augmentation of dendritic excitability could help explain how cortically-evoked synaptic activity is biased towards the direct pathway in SAPAP3 KO mice (Ade et al., 2016).

Why does deletion of a synaptic gene cause cell-wide changes in excitability? One possibility is that the changes in excitability represent homeostatic adaptations to impaired excitatory drive (Fieblinger et al., 2014). Another (non-mutually exclusive) possibility is that normal neuromodulatory processes are altered. ACh, via muscarinic receptors, modulates K^+^ and voltage gated Ca^2+^ that regulate SPN somatic and dendritic excitability (Goldberg et al., 2012; Oldenburg and Ding, 2011). Though we report that striatal ACh release and muscarinic signaling are altered in SAPAP3 cKI^-/-^ mice, however, blockade of muscarinic receptors did not normalize dSPN dendritic excitability. Similarly, while pathological dopamine signaling has been observed in OCD patients and mouse models of OCD (Bellini et al., 2018; Fineberg et al., 2011; Pauls et al., 2014; Wood et al., 2018), the fact that dendritic excitability was unchanged in D2 dopamine receptor expressing iSPNs makes altered dopaminergic modulation an unlikely culprit in this instance (Day et al., 2008; Fieblinger et al., 2014). One other possibility is that changes in excitability may involve pathologically elevated mGluR5 signaling (Chen et al., 2011; Wan et al., 2011). Indeed, mGluR5 receptors can modulate somatic excitability and SPN dendritic Ca^2+^ signaling (D’Ascenzo et al., 2009; Plotkin et al., 2013), and blockade of mGluR5 receptors normalizes imbalanced cortically-driven striatal output (Ade et al., 2016). It is unclear how elevated mGluR5 signaling might lead to pathway-specific changes in dendritic excitability though, given that both dSPNs and iSPNs are hyper-responsive to mGluR5 activation in SAPAP3 KO mice (Chen et al., 2011; Wan et al., 2011).

Strikingly, constitutive deletion of SAPAP3 led to an increase in dorsal striatum CIN density and evoked ACh release. This is intriguing in light of two lines of evidence. First, recent studies have discovered that children with Pediatric Autoimmune Neuropsychiatric Disorder Associated with Streptococcus (PANDAS), a condition associated with early onset OCD symptoms, exhibit IgG antibodies that bind to and decrease the activity of CINs (Frick et al., 2018; Xu et al., 2021). Second, both ablation of striatal CINs and augmentation of striatal ACh release (by overexpression of the vesicular ACh transporter) accentuate stress- and/or psychostimulant-evoked stereotypies in mice, with CIN ablation leading to tic-like stereotypies or extending cocaine-induced stereotypies and elevated ACh release exacerbating psychostimulant-induced stereotypies (Aliane et al., 2011; Crittenden et al., 2014; Xu et al., 2015). Why would both elevations and presumed decreases in striatal ACh release enhance behavioral stereotypies? One possibility is that there’s simply an optimal level of ACh release for normal striatal function, with deviations too far above or below leading to pathological behavior. But the answer may be more nuanced, as implied by our observations of both hyper- and hypo-cholinergic pathologies in SAPAP3 cKI^-/-^ mice: while evoked ACh release is elevated, signaling through muscarinic receptors is attenuated in SPNs. The potential impact of hypercholinergic release will be discussed below, but impaired muscarinic signaling in SAPAP3 cKI^-/-^ SPNs is consistent with the observation that muscarinic antagonists can prolong psychostimulant-induced stereotypies (Aliane et al., 2011).

The complex and often opposing consequences of CIN function on repetitive behaviors are likely due to the rich milieu of muscarinic and nicotinic ACh receptors located throughout the striatum. While muscarinic signaling in SPNs is impaired, other receptors (or downstream signaling arms of the same receptors) may track changes in ACh very differently, even in pathway-specific ways (Abudukeyoumu et al., 2019; Assous, 2021; Goldberg et al., 2012). Of particular relevance to the compulsive behaviors at the center of this study are the impacts of ACh on dopamine release. Activation of presynaptic nicotinic ACh receptors on nigrostriatal axon terminals triggers the release of dopamine (Rice and Cragg, 2004; Threlfell et al., 2012). It’s plausible that the increased density of striatal CINs and augmented ACh release in SAPAP3 cKI^-/-^ mice will enhance nicotinic ACh receptor mediated dopamine transients, a process that would ultimately promote cortical engagement of the direct pathway (Gerfen and Surmeier, 2011).

Besides membrane excitability and Ca^2+^ flux, muscarinic receptors also directly modulate NMDA receptor function: M1 type muscarinic receptors can enhance NMDA receptor currents (Calabresi et al., 1998), while M4 type receptors can suppress them (Shen et al., 2015). Importantly, all SPNs express M1 receptors but M4 receptors are preferentially found in dSPNs (Hersch et al., 1994; Shen et al., 2015). Future studies are necessary to determine if M1/4 receptor modulation of NMDA receptors is attenuated or enhanced by elevated ACh release in SAPAP3 cKI^-/-^ mice. NMDA receptor expression and function are profoundly altered in SAPAP3 KO mice (Corbit et al., 2019; Hadjas et al., 2020; Welch et al., 2007), a pathology that may be even further abrogated by aberrant cholinergic modulation. Adding to the appeal of this possibility, we observed that although basal muscarinic modulation of individual dendritic spines was globally attenuated in iSPNs of SAPAP3 cKI^-/-^ mice, a significant percentage of spines in dSPNs actually became more excitable after muscarinic blockade. Given the cell-type specific differences in M1 and M4 expression noted above, one interpretation is that M4 signaling is intact in SAPAP3 cKI^-/-^ mice, setting the stage for pathological pathway- and synapse-specific modulation of excitatory inputs by ACh.

### Limitations of the study

This study describes opposing alterations in cholinergic signaling in the SAPAP3 cKI^-/-^ mouse model of OCD: 1) increased striatal CIN density and evoked ACh release and 2) decreased basal ACh signaling through muscarinic receptors. How striatal CIN density is enhanced is not clear, though we note that CINs are not the only class of striatal interneurons whose numbers are impacted by deletion of SAPAP3 (Burguiere et al., 2013). While CINs are the major source of ACh in the dorsal striatum, and they are even more abundant after the loss of SAPAP3, they are not the only source. It remains to be determined if ACh release from other sources, such as the pedunculopontine nucleus (Dautan et al., 2020), is augmented as well. Finally, the focus of this study is on the *basal* changes in excitability and modulation of SPNs that emerge in a mouse model of OCD. While basal muscarinic signaling in SPNs is impaired in SAPAP3 cKI^-/-^ mice, limitations in ACh sensor sensitivity under our experimental conditions preclude the ability to accurately estimate *basal* levels of ACh in *ex vivo* brain slices from WT and mutant mice.

## Methods

### Animal subjects

All experimental procedures were performed in accordance with the United States Public Health Service *Guide for Care and Use of Laboratory Animals* (Institute of Laboratory Animal Resources, National Research Council), were approved by the Institutional Animal Care and Use Committee at Stony Brook University. Animals were housed 2-5 per cage in an environmentally controlled room (20-23°C, 30%-70 humidity, 12h light/dark cycles) with food and water available *ad libitum*, and cages were cleaned once per week. Due to tendency of mutant animals to develop facial and bodily lesions, enhanced animal welfare care was deemed necessary and topical treatments with triple antibiotic ointment were administered by veterinary staff as needed. All experiments were performed in 3-to 7-month-old male and female C56BL/6 mice crossed between a SAPAP3 null line (conditional knock-in, generated by Dr. Qiangge Zhang and Dr. Guoping Feng) and 1 of 2 BAC transgenic lines that express fluorescent reporters either under the D1 (drd1-tdTomato) or D2 (drd2-eGFP) dopamine receptor promoter (Gong et al., 2003; Shuen et al., 2008). Experiments performed on SAPAP3^-/-^ mice were carried out after the onset of compulsive grooming and matched alongside wildtype controls of a comparable age.

### Behavioral Analyses

For the assessment and quantification of grooming behavior, experimental subjects were habituated to a 13cm diameter acrylic cylinder for 10 mins daily, 3 days prior to video acquisition for a duration of 10 minutes on the fourth day. Behavioral tests were carried out no more than 24 hours prior to experimentation and grooming behavior was manually analyzed, as per the grooming bout parameters described in Welch et al. 2007, to confirm the overgrooming phenotype in SAPAP3 cKI^-/-^ mice. (see **Figure S1**).

### Ex vivo Slice Electrophysiology

Acute parasagittal slices (275μm) containing the dorsolateral striatum were obtained from experimental subjects following anesthetization with ketamine/xylazine (100mg/kg/7mg/kg) and transcardial perfusion with ice-cold artificial cerebral spinal fluid (ACSF) containing (in mM): 124 NaCl, 3 KCl, 1 CaCl_2_, 1.5 MgCl_2_, 26 NaHCO_3_, 1 NaH_2_PO_4_, and 14 glucose, continuously bubbled with carbogen (95% O_2_ and 5% CO_2_). Slices were cut using a VT-1000 S vibratome (Leica Microsystems, Buffalo Grove, IL) and transferred to a holding chamber where they were incubated at 32°C for 45 mins in ASCF containing (in mM) 2 CaCl_2_ and 1 MgCl_2_, after which they were removed and acclimated to room temperature (~21°C) for 15 mins before recording. Using a Bruker 2-photon imaging system with integrated electrophysiological capabilities, SPNs were identified via tdTomato or eGFP fluorescence and patch-clamped for whole-cell recording (3.5-5MΩ electrodes). Current clamp recordings were performed with an internal solution containing (in mM): 135 KMeSO_4_, 5 KCl, 10 HEPES, 2 ATP-Mg^2+^, 0.5 GTP-Na^+^, 5 phosphocreatine-tris, 5 phosphocreatine-Na^+^, 0.1 Fluo-4 pentapotassium salt, and 0.05 Alexa Fluor 568 hydrazide Na^+^ salt. Fluo4 precluded the need for EGTA in Ca^2+^ imaging experiments and Alexa 568 was used to visualize cell bodies, dendrites, and spines. Electrophysiological recordings were digitally sampled at 30kHz and filtered at 1kHz using a Multiclamp 700B amplifier (Molecular Devices, San Jose, CA).

### 2-photon Ca^2+^ laser scanning microscopy, Ca^2+^ imaging and ACh sensor measurements

dSPNs and iSPNs were identified via somatic tdTomato and enhanced-GFP 2-photon excited fluorescence using an Ultima Laser Scanning Microscope System (Bruker Nano, Inc., Middleton, WI). For simultaneous electrophysiological & 2-photon Ca^2+^ imaging experiments, green and red fluorescent signals were obtained using 810nm pulsed light excitation (90MHz) (Chameleon Ultra II, Coherent, Inc., Santa Clara, CA). Patched SPNs were allowed to equilibrate to the internal solution prior to any experimental Ca^2+^ imaging for a minimum of 15 minutes after patch rupture to ensure adequate filling of distal dendrites. *Ca^2+^ imaging:* Dendritic Ca^2+^ transients were measured using Fluo4, as previously described (Day 2008). Line scans of green (G) and red (R) fluorescence were acquired at proximal (45μm-60μm) and distal (90μm-130μm) adjacent dendritic spines and shafts at 5.342ms and 512 pixels per line with 0.0776μm^2^ pixels at 10μs dwell. Ca^2+^ transients were expressed as ΔG/R, and dendritic indexes were quantified as distal Ca^2+^ transient area (ΔG/R•ms) divided by the largest proximal Ca^2+^ transient area (for single bAPs) or the average of 2-5 proximal Ca^2+^ transient areas (for bAP bursts), all acquired from dendritic shafts (Carrillo-Reid et al., 2019). Single bAPs were generated by a somatic current injection (2nA, 2ms) and bursts of bAPs were delivered at 5Hz, each burst containing 3 bAPs at 50Hz (20ms interval). Line scans began 300ms before stimuli and continued for 698ms (for single bAPs) or 1670ms (for bAP theta bursts) after the stimuli ended. All drugs were dissolving in the external ACSF and bath applied for a minimum of 10 minutes prior to comparative experimentation. *ACh sensor measurements:* To evoke ACh release, a concentric stimulating electrode powered by an ISO-Flex Stimulus Isolator (both from Microprobes for Life Science) was centered on the y-plane and placed just outside of the objective field. Spiral line scans (920nm, 21.2ms with 0.0776μm^2^ pixels and 10μs dwell) were placed over infected neuropil and acquired sequentially, at least 5 seconds apart, while delivering single 1ms electrical stimuli of increasing intensities. Line scans began 1000ms before stimuli and continued for 4000ms after the stimuli ended.

### Surgical procedures

All mice subjected to surgical procedures were induced and maintained under anesthesia via inhalation of vaporized isoflurane (3% for induction and 1%-2% for maintenance during surgery) and administered a post-operative analgesic, Meloxicam (5 mg/kg), via subcutaneous injection as needed. Prior to any surgical procedure, complete sedation was confirmed via tail pinch, animals were placed in a stereotaxic apparatus, the scalp was shaved and disinfected, and eyes were covered with an eye-protective gel (Puralube Ophthalmic Ointment, Dechra Veterinary Products). A 1μl Hamilton Neuros micro-volume syringe was used for unilateral dorsolateral striatum injections (−2.20mm ML, 0.62mm AP, −2.80mm DV) and manipulated via a motorized stereotaxic injector (Quintessential Stereotaxic Injector QSI^™^, Stoelting Co.) controlled by Angle Two software (Leica Biosystems). 200nl of AAV5-hSyn-ACh3.0(ACh.4.3) (ACh3.0; titer: ≥1•10^13^vg/ml) (Jing et al., 2020) were injected over 5 minutes and a further 5 minutes was allowed for diffusion before slowly withdrawing the needle. All subjects were monitored daily for pain for 72h and sacrificed for experimentation after a postoperative period of 3-4 weeks.

### Immunohistochemistry

Experimental subjects were anesthetized by isoflurane inhalation before transcardial perfusion with PBS, then 4% paraformaldehyde (PFA) (Electron Microscopy Sciences) before brain extraction (note that all IHC dilutions are in 1x phosphate-buffered saline [PBS]). Extracted brains were fixed in 4% PFA for a further 12 hours at 4°C, then cryopreserved with 30% sucrose in 1x PBS for 48 hours at 4°C. Fixed brain tissue was frozen in O.C.T. (Fisher) and sliced coronally at 50μm thickness using a cryostat (Leica). For antibody labeling, brain slices were blocked for 2h at RT (3% normal donkey serum [Sigma D9663], 0.1% triton-x), stained with 1’ ab for 12h at 4°C (EMD Millipore [ab144P, RRID: AB_2079751] goat anti-ChAT, 1:500 dilution), washed 3 x 10 minutes (0.5% NDS, 0.1% triton-x), stained with 2’ ab for 2h at RT (Abcam [ab150132, RRID: AB_2810222] donkey anti-goat Alexa Fluor 594, 1:2000 concentration), then washed 3 x 10 minutes as above before placing in PBS and mounting onto microscope slides with Fluoromount-G (SouthernBiotech). Another antibody combination was used alongside ChAT labelling (rabbit anti-uOR [Abcam [ab10275, RRID: AB_2156356]) 1’ and donkey anti-rabbit 488 (Invitrogen [A-21206, RRID: AB_2535792] 2’), but not analyzed as part of this study. Coronal sections were collected and binned at 200μm increments from the rostral striatum (1.25mm-1.05mm bregma) to the caudal striatum (0.45mm-0.25mm bregma) portions of the pre-commissural (anterior) striatum.

### Immuno image acquisition & analysis

Images were obtained using a VS120-S6-W virtual slide microscope (Olympus) at 20x magnification. Images were composed of 30 1μm z-planes. Image analysis was performed by two blinded investigators using ImageJ (Fiji [RRID: SCR_002285]). The 15 most in-focus planes in the z-stacks were max-projected and thresholded. Semi-automated quantification of cell bodies was performed using ImageJ’s Particle Analysis function. Identified cells were manually checked for accuracy. Striatal regions were manually outlined using the lateral ventricles, callosum, and a straight line from the inferior ventricular apex to the inferomedial corner of the piriform cortex to reproducibly demarcate the striatum. Dorsal and ventral striatal domains were established by a horizontal partition dividing the entire striatum into two equal areas.

### Quantification and statistical analyses

Electrophysiological recordings and Ca^2+^ transients were analyzed using Igor Pro (WaveMetrics, Inc., Lake Oswego, OR [RRID: SCR_000325]). Statistical differences were examined using 2-sided Mann-Whitney U, Wilcoxon signed-rank (for paired comparisons), 2-way ANOVA (for grouped comparisons), or Fisher’s exact. Differences were considered significant if p<0.05. All statistical tests were performed in GraphPad Prism v9 (GraphPad Software, La Jolla, CA [RRID: SCR_002798]) prior to and following a Grubb’s test for outliers (alpha= 0.05), also in Prism.

## Acknowledgements

We thank W. Akmentin and Jennifer Wilking for technical support. This study was supported by NIH NINDS R01 NS104089 to J.L.P.

## Author contributions

Conceptualization: J.M.M. and J.L.P; Investigation and analysis: J.M.M., A.B., K.J., Z.B.H. and E.M.P.; Supervision and generation of SAPAP3 cKI^-/-^ mice: Q.Z. and G.F.; Writing: J.M.M. and J.L.P.; Supervision and analysis: J.L.P.

## Supplemental information

### Generation of SAPAP3 cKI^-/-^ mice

Sapap3 conditional knock-in (cKI) mice were generated using a similar strategy described previously (Zhang et al., 2016; Mei et al., 2016). In brief, three exons (Ensemble ID: ENSMUSE00001223455, ENSMUSE00000400411, ENSMUSE00000335250) and the introns between those exons of the Sapap3 gene were inverted and cloned into the FLEX targeting vector (Mei et al., 2016). Mouse R1 ES cells were used to generate the clones that contain the correct targeting allele. One of those ES clones was implanted into C57 blastocysts to produce the chimeric founder, which was crossed with betaActin-FLP mice (The Jackson Laboratory, stock #005703) to generate the Sapap3 cKI mouse line. Those mice were backcrossed to C57BL/6J (The Jackson Laboratory, stock #000664) for >5 generations before use in experiments. In the absence of Cre recombinase, the inverted exons and introns would introduce reading frame shift and cause degradation of Sapap3 transcripts.

**Supplemental Figure 1:**
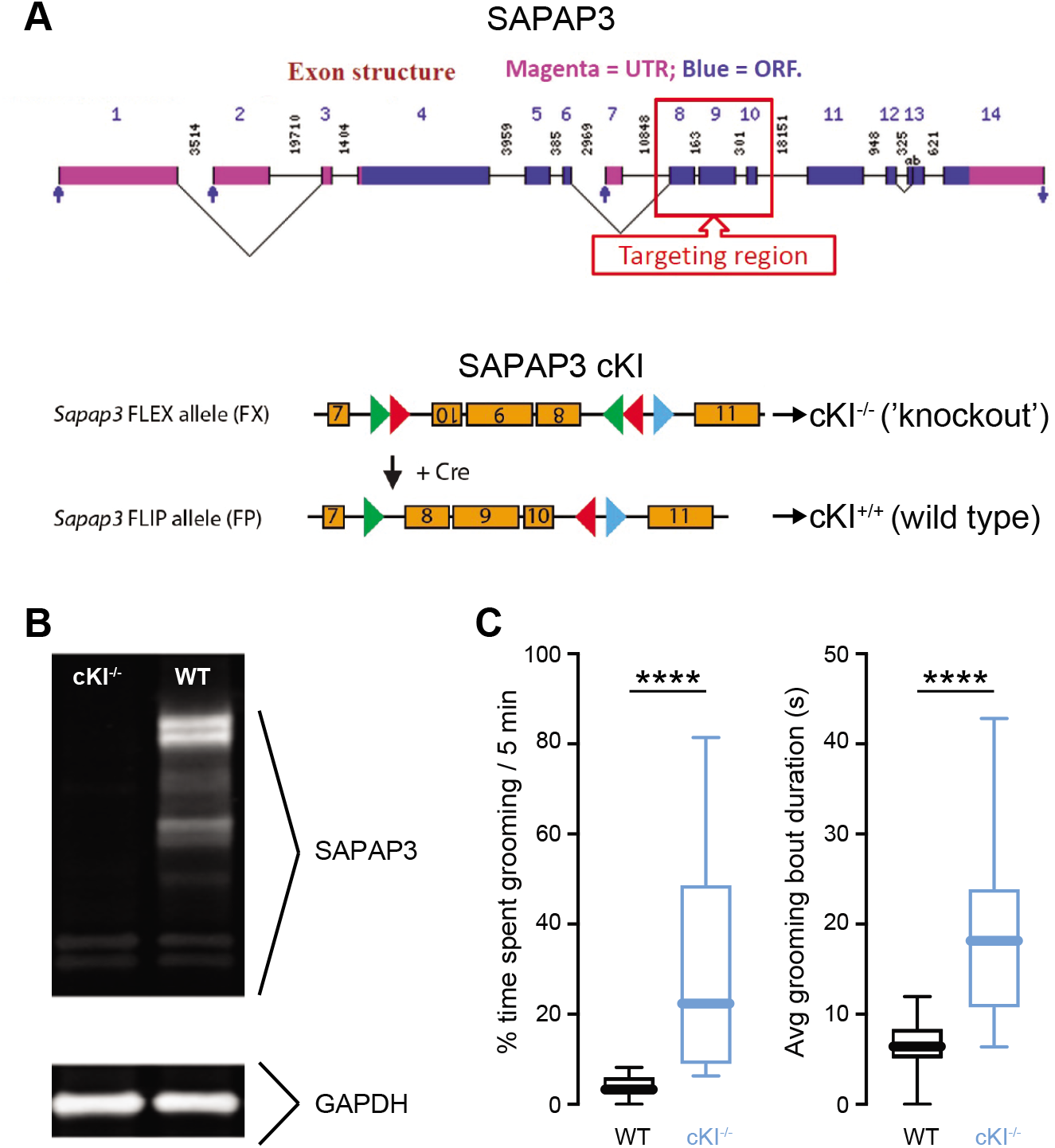
Generation and validation of SAPAP3 cKI^-/-^ mice. (A) (Top) Exon structure of the mouse SAPAP3 gene, indicated the region targeted to create the cKI line. From NIA mouse gene index. (Bottom) Strategy for creating cKI^-/-^ mice. (B) Validation that SAPAP3 protein is lost in Sapap3 cKI^-/-^ mice. Lanes are brain lysate from WT (left) and SAPAP3 cKI^-/-^ (right) mice, probed with rabbit anti-SAPAP3 (top) and mouse anti-GAPDH (bottom). (C) Box plots showing percentage of time WT (black) and SAPAP3 cKI^-/-^ (blue) mice spent grooming (left) and average grooming bout durations (right) within a 5-minute timeframe. ****p<0.0001. Mann-Whitney, p<0.0001, WT n=18, SAPAP3 cKI^-/-^ n=25.

**Supplemental Figure 2:**
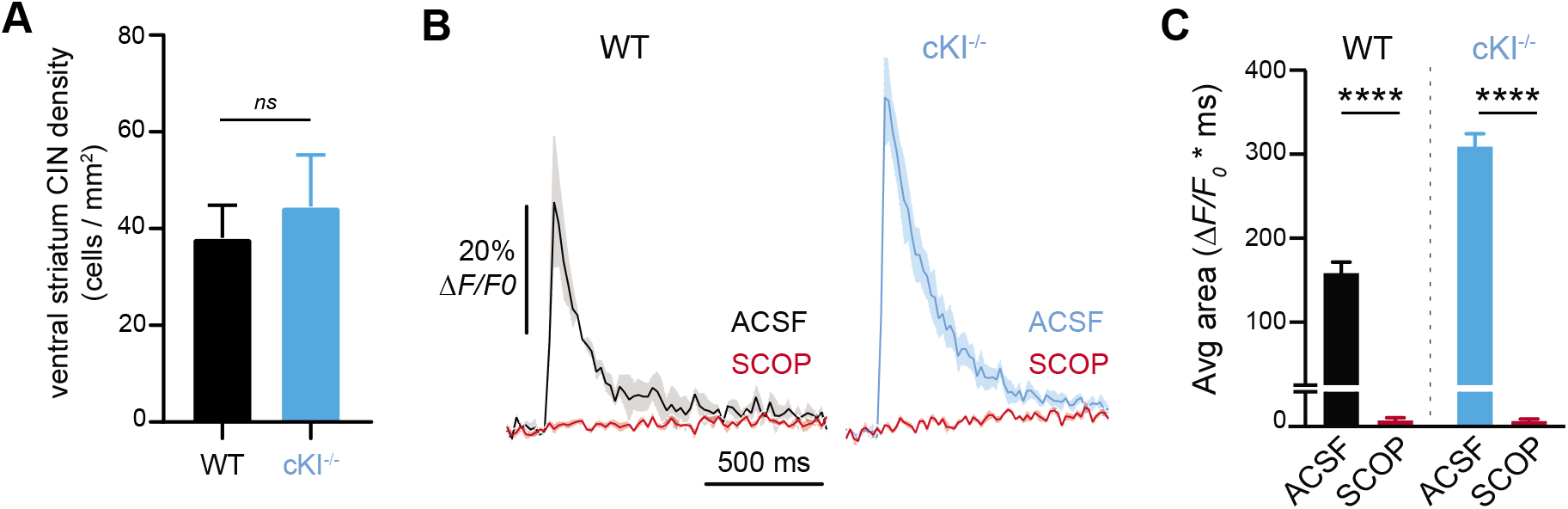
Ventral striatum CIN densities and ACh3.0 sensor specificity. (A) Box plots showing CIN density in the ventral striatum of WT (black) and SAPAP3 cKI^-/-^ (blue) mice. (B) Averaged line scans measuring changes in ACh3.0 fluorescence in dorsal striatum neuropil in response to a 3mA electrical stimulus in WT (black) and SAPAP3 cKI^-/-^ (blue) mice before (ACSF) and after a 10μM scopolamine hydrobromide (SCOP). Shaded areas are S.E.M. (WT n=8, SAPAP3 cKI^-/-^ n=10). (C) Bar graphs showing average changes in ACh3.0 fluorescence in WT (black) and KO (blue) mice before and after SCOP. Measurements are in response to 3mA electrical stimuli. (WT n=8, SAPAP3 cKI^-/-^ n=10). ****p<0.0001.

